# Reproduction emerges from ecological interactions at the onset of multicellularity

**DOI:** 10.1101/2025.10.08.681199

**Authors:** Alexandre P. Fernandes, Renske M. A. Vroomans, Enrico Sandro Colizzi

## Abstract

At the origin of multicellularity, genetic programs used by single-celled organisms to interact with their environment become organised into a developmental program. While previous work has clarified the selective advantages of simple multicellularity, the origin of development remains unclear. Here, we investigate the ecological origin of a fundamental developmental process — multicellular reproduction — using a computational model where cells forage in a structured environment, and evolve adhesion and regulated environmental responses. Multicellular modes of reproduction are not pre-specified in the model, but must emerge from the evolving dynamics under ecological selection. We find that distinct multicellular life cycles evolve depending on the spatial distribution of resources. Among these are life cycles with a single-celled propagule phase the most prevalent reproductive strategy in multicellular life that evolves as a dispersal strategy to reach new resources. These propagules form through the activation of molecular programs co-opted from the unicellular ancestor, where they mediated interactions with neighbours in its ecological context. Once propagules evolve, multicellular lineages can invade environments previously dominated by unicellular competitors, showing this strategy is adaptive beyond the conditions that permit its evolution. Altogether, our results show that developmental regulation evolves through co-option of ecological interactions during the transition to multicellularity.

**Significance statement:** Reproduction is a universal feature of life. Yet, the evolution of multicellularity transformed it fundamentally: while single-celled organisms reproduce via cell division, reproduction in multicellular organisms is a complex process involving the coordination of many cells. How these new forms of multicellular reproduction first evolved is currently unknown. Using a computational model, we study how group reproduction emerges from the collective dynamics of individual cells. The model shows that unicellular ancestral life cycles can be repurposed as propagules used for reproduction in multicellular species, suggesting that genetic co-option is a key mechanism through which early development evolves.

## Introduction

While unicellular organisms reproduce through cell division, multicellular reproduction is a collective process that requires coordination of cell division, collective behaviour, and spatial organisation to generate a new individual. Modern multicellular organisms have evolved a diverse array of reproductive strategies, which can be broadly divided into two main categories (1, 2): (i) unicellular propagation, where the organism reproduces by generating single-celled reproductive units, and (ii) multicellular propagation, where the organism divides into two multicellular groups of equal or unequal size. Unicellular propagation is abundant across the tree of life, including sexual reproduction through gametes and formation of unicellular vegetative propagules or spores (3, 4). Although not as common as unicellular propagation, multicellular propagation occurs in all major multicellular clades, including bacteria (5, 6), algae (7), fungi (8), plants (9, 10), and animals (11, 12) (see (13, 14) for comprehensive reviews of multicellular life cycles). Additionally, both forms of reproduction have been observed to evolve in wet-lab experiments (15–18).

But what drives the evolution of different reproductive strategies? Computational models have suggested that unicellular propagules reduce genetic conflicts between cells within the multicellular organism (19, 20) or slow down the accumulation of mutations (1), while experimental evolution supports their role in enabling higher-level individuality (21). Conversely, multicellular reproduction can be advantageous in cell clusters with rigid cell connections that limit cell division, enabling cell growth in regions previously constrained by the cluster’s geometry (22). More broadly, the selective advantage of unicellular or multicellular replication may depend on how cell birth and death rates scale with group size (23) and on the effect of cooperation between cells in multicellular groups (24).

The ecological context also plays an important role in shaping the evolution of multicellular reproduction. Many organisms have evolved unicellular propagules, such as spores, pollen, or cysts, that can disperse passively via wind or water and are often more resistant to environmental stress than their multicellular progenitors. This makes them well-suited for survival in harsh environments and an effective means of dispersal. Models and experiments suggest that diverse life cycles, including those alternating between unicellular and multicellular stages, can evolve in response to fluctuating environments (25–28), or arise through self-organisation of multicellular groups that can reproduce autonomously due to adhesion interactions between cells (29). The predicted influence of the environment on the evolution of multicellular reproduction is also evident in nature, where seasonal fluctuations in nutrient availability regulate life cycle transitions in many eukaryotes. For example, nutrient scarcity triggers the multicellular phase in the life cycle of the slime mould *Dictyostelium discoideum*, whereas the Holomycota *Fonticula alba* forms collective feeding streams only when bacterial food is available (30). Conversely, the emerging multicellular group can itself alter the ecological landscape, affecting the evolutionary trajectories of cellular properties (31) and creating new ecological niches (32).

The ecological and genetic contexts have also played a role in the origin of multicellularity itself. As cells formed tighter associations during the transition to multicellularity, new ecological functions — e.g. predation avoidance (15, 18) or enhanced resource capture (33, 34) — selected for traits that increased group sizes, improved the spatial integrity of the cluster, and mediated collective responses to environmental fluctuations (1, 35, 36). Comparative phylogenetic studies show that these selective demands at the onset of multicellularity were largely met by coopting the pre-existing molecular toolkit (37, 38): proteins used for adhesion with the substrate or sensing environmental cues were repurposed for itercellular adhesion (39, 40), while cell-cycle and stress regulators became involved in cellular differentiation (41, 42). However, the contribution of the co-option of these ancestral modules to the evolution of development and reproduction at the onset of multicellularity remains unknown. Promising experimental results show that reproductive propagules can readily evolve after multicellularity, suggesting that they could have evolved via co-option of an ancestral unicellular life cycle (17).

To clarify how ecological pressures influence the spatial organisation of cells during reproduction and how the co-option of ancestral modules enables novel ecological functions, we need a modelling framework that integrates the evolution of gene regulation with ecological interactions between cells and their environment (43, 44). Existing models, however, typically lack a spatially-explicit environment, abstract away genetic regulation of reproductive behaviour, or restrict the evolution of reproductive strategies to a few, pre-selected options. To address this issue, we used a bottom-up approach to study the evolution of multicellular reproduction. We developed a model where a population of cells inhabits a structured environment and consumes resources to survive and replicate. We model a minimal set of cellular traits — intercellular adhesion, chemotaxis, and cell division — that are widespread across the eukaryotic tree of life (4). With this setup, we investigate how spatially structured ecological interactions among unicellular individuals give rise to reproduction at the onset of multicellularity. We characterise these developmental dynamics and show that they evolve by co-opting pre-existing life-cycles and are stabilised through an eco-evolutionary feedback.

## Results

### Model setup

To investigate the evolution of reproductive strategies in nascent multicellular organisms, we constructed a hybrid Cellular Potts Model (45–47) where a population of cells forages for food on two-dimensional space (Fig. 1A). Food is introduced at a constant rate into the environment as circular “patches”, each composed of several food units. The food patches release a chemoattractant, creating a gradient around them that cells can follow to locate and consume resources, which are stored as metabolic reserves. If all the food in a food patch is consumed, it stops generating the chemoattractant signal, allowing cells to re-route to the next source of food. Cells have a constant metabolic rate that depletes their metabolic reserves. When the cell’s reserves are exhausted, it dies and is removed from the lattice. We also include a small uniform probability of death due to other factors which are not modelled explicitly, such as metabolic stress.

**Fig. 1.**
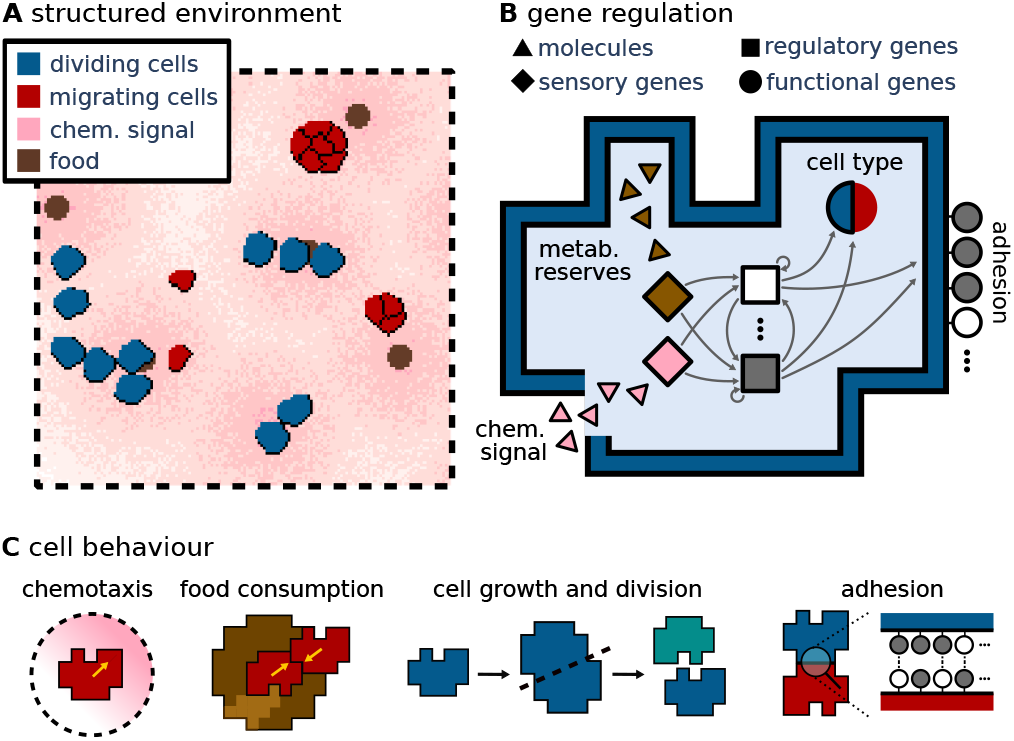
Model setup. **A** Cells live in a structured environment where they forage for resources by following the chemoattractant signal that diffuses from patches of food. Here, a small section of the 2000×2000 lattice is shown. **B** Diagram of a cell’s gene regulation. Cell behaviour is controlled by a Boolean GRN that senses the amount of metabolic reserves accumulated in the cell and the strength of the chemoattractant signal. These inputs are processed by a layer of 18 regulatory genes that will determine the cell state (migrating or dividing) and the activation state of the 16 adhesion genes. Active genes are shown in grey, inactive in white. Arrows show the connections between genes in the GRN. **C** We model two cell states: migrating and dividing. Cells can accumulate metabolic reserves by consuming food if they share a lattice site with a food site and adhere to their neighbours when they express complementary combinations of adhesion proteins.

Cellular behaviour is controlled by a Gene Regulatory Network (GRN) composed of sensory, regulatory, and functional genes (Fig. 1B). The sensory genes receive as input the concentration of the chemical signal at the cell’s location and the amount of internal metabolic reserves. These inputs are then processed by a layer of regulatory genes that control the activation of the downstream functional genes. Functional genes dictate cell behaviour. We model a minimal set of cellular behaviours commonly observed in microbial eukaryotes (48–51): cells can migrate following the chemical signal diffusing from food sources, consume food from the environment, divide, and adhere to their neighbours (Fig. 1C). The cell state — either migrating or dividing — is determined by the output of the first functional gene. We model these two behaviours as mutually exclusive, following the observation that migration and division require conflicting use of the cytoskeleton in many eukaryotes (52–54). Migrating cells perform chemotaxis by biasing their movement towards the direction of the perceived chemical gradient (55), while dividing cells cannot actively migrate and grow over a fixed period before dividing into two daughter cells. When a cell divides, the parameters of the GRN of one daughter cell can mutate with a small probability, while the other cell inherits the parental parameters. Both daughter cells inherit half of the metabolic reserves of their mother. An additional set of sixteen functional genes encodes adhesion proteins that are expressed on the cell membrane and mediate adhesion between adjacent cells. These proteins are modelled as ligands and receptors, and adhesion strength (*γ*) between adjacent cells increases with the degree of complementarity between their ligands and receptors (for all the model details, see SI Appendix, Model description).

Together, these components create a structured dynamic environment in which individual cells must integrate internal and ecological cues to survive. The mutations to the GRN during cell division introduce genetic variation into the population. This variation, combined with differential survival and reproduction under ecological constraints, drives an evolutionary process without directly specifying its outcome, which is thus free to emerge spontaneously. In the following sections, we examine how this setup gives rise to diverse life cycles that can coordinate multicellular reproduction.

### Evolution of diverse life cycles is mediated by food distribution

Using this setup, we ran 72 simulations each starting with a population of 800 cells. The cells start with a randomised GRN which can evolve throughout the simulation. Simulations ran for 10^8^ time steps in one of six environmental conditions (EC 1-6) that differ in the heterogeneity of food distribution. We explore food heterogeneity systematically across ECs, by doubling food patch area and halving creation frequency between successive ECs, while holding total food influx constant (Table S1 and Fig. S1). EC 1 has a relatively even resource distribution, with many small, closely spaced food patches, whereas in EC 6 resources are concentrated into a few large, distant patches. At the end of each simulation, we characterised the dominant evolutionary outcome by extracting the last common ancestor of the population and analysing its behaviour through follow-up simulations without mutations (see Materials and methods, Non-evolutionary simulations).

Three main life cycles evolved across our simulations: a strictly unicellular life cycle, a strictly multicellular life cycle, and a mixed life cycle that is multicellular but regulates the formation of unicellular propagules (Fig. 2A and Videos S1-4, see Materials and methods, Categorisation of life cycles). In all cases, cells have evolved to migrate up the chemoattracting gradient when their metabolic reserves are low, and divide after consuming food from the environment. In the multicellular + unicellular propagules life cycle (Fig. 2A, blue), cells adhere to form multicellular clusters during migration, but stop adhering once they initiate division. These dividing cells are left behind as unicellular propagules, while the multicellular cluster continues migration following a new gradient after local resources are depleted. After dividing, the daughter cells of the unicellular propagule resume migration and re-activate adhesion genes, forming a newborn multicellular cluster that follows the chemoattracting gradient to find new resources, thus completing the life cycle.

**Fig. 2.**
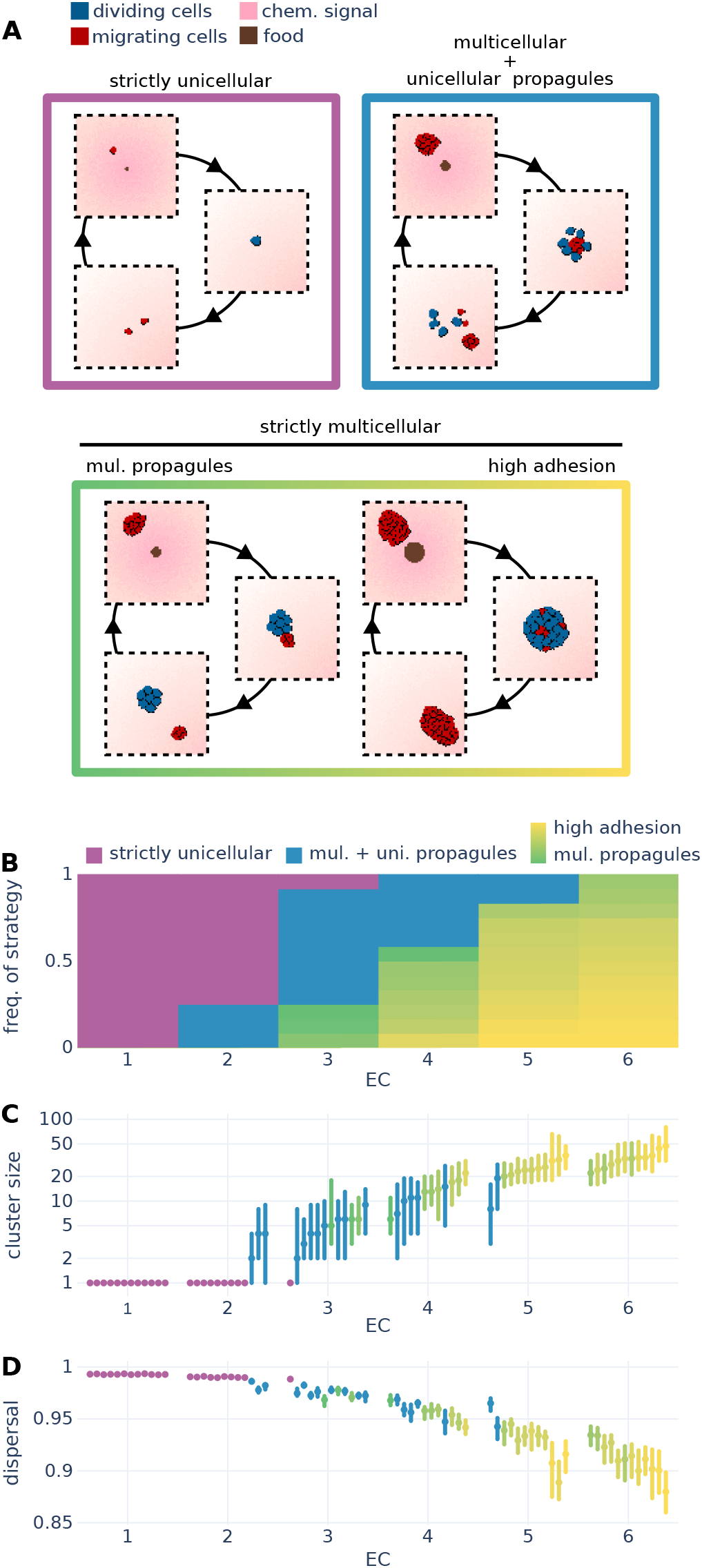
Spatial heterogeneity of resources drives the evolution of complex multicellular life cycles. **A** Three categories of life cycles evolve in our simulations: strictly unicellular, multicellular + unicellular propagules, and strictly multicellular. Strictly multicellular strategies form clusters of dividing cells whose adhesion to the migrating group ranges from loose multicellular propagules that readily detach to highly adhesive groups that rarely split. **B** Distribution of evolved life cycles in different environmental conditions. **C** Cluster sizes and **D** dispersal of all evolved lineages (median and interquartile ranges, see Materials and methods, Measure of dispersal). Lineages were ordered by median cluster size.

Some strictly multicellular lineages also form propagules. In this case, the propagule is multicellular and is formed by dividing cells that downregulate adhesion to migrating cells, while still adhering to each other. We observe a spectrum of multicellular reproductive stratgies: at one end are lineages that genetically encode the formation of propagules by consistently downregulating adhesion between migrating and dividing cells (green in Fig. 2), while at the other end are lineages whose clusters divide rarely, and exclusively by phyisically tearing themselves apart when different parts of the cluster are attracted by separate food patches (yellow in Fig. 2).

Food heterogeneity strongly impacts the evolution of reproductive strategies in the model (Fig. 2B). Strictly unicellular strategies evolve most often when food is homogeneously distributed (EC 1-2), while high-adhesion multicellular strategies usually evolve when food distribution is heterogeneous (EC 5-6). Intermediate environments are characterised by the dominance of unicellular and multicellular propagules (EC 3-4). Lineages with different life cycles show contrasting adaptive functional characteristics that aid survival in the environments where they evolved. For example, high-adhesion multicellular lineages have larger cluster sizes when compared to strictly unicellular and propagule-forming lineages, since their cells are concentrated in a few clusters that rarely divide (Fig. 2C). These large multicellular clusters display collective chemotaxis, allowing them to migrate faster than small clusters or single cells. This happens because cells in a group can integrate spatial information about the patchy chemoattractant signal by pulling and pushing neighbouring cells, allowing them to correct each other’s trajectories towards the source of food (56). Collective migration, and thus stable multicellularity, becomes more advantageous the longer the distance cells must migrate to reach food, which explains why high-adhesion multicellular life cycles dominate the most heterogeneous environmental conditions (see SI Appendix, Selective advantages of multicellularity and unicellularity, Video S5, Fig. S2A and B).

Conversely, unicellular lineages disperse better through the environment when compared to multicellular lineages (Fig. 2D). Separating after division helps cells colonise and consume multiple food patches at once, which is especially advantageous when resources are split among many food patches in the most homogeneous environments (see SI Appendix, Selective advantages of multicellularity and unicellularity, Video S6, Fig. S2C and D).

Previous studies suggested that reproduction through single-celled units could have evolved as a mechanism to reduce genetic conflict in populations of clonal multicellular organisms (1, 19). To determine whether this genetic factor could explain the evolution of single-celled propagules in our model, we analysed the phylogeny of evolved cell populations from the simulations (Fig. S3). We found that the cell clusters of both strictly multicellular and propagule-forming populations are highly heterogeneous due to their aggregative nature, indicating that in this case the reduction of genetic conflicts does not play a role in the evolution of propagules. Instead, the main factor driving the evolution of reproduction via propagules seems to be ecological. In the intermediate environments where unicellular and multicellular propagules evolve, food patches are far apart enough that collective migration is advantageous, while being plentiful enough that dispersal is critical for patch colonisation. As a result, propagule-forming lineages have evolved a multicellular migrating phase that allows them to form clusters to benefit from group migration (Fig. 2C), together with the ability to make unicellular or multicellular propagules for reproduction and dispersal (Fig. 2D).

### Development of reproductive strategies: evolved co-ordination of adhesion and cell state regulation

Because the formation of propagules detaching from migrating clusters emerged spontaneously in our model, we next characterise the regulatory mechanisms that make it possible. We started by quantifying adhesion strength between cells expressing different cell states across our simulations. This shows that propagule-formers coordinate the expression of their cell state and adhesion genes, leading to migrating cells that adhere strongly to other migrating cells but not to dividing cells (Fig. 3A, blue and green). The gene regulatory network achieves this coordination by evolving regulatory genes that co-regulate cell state and the expression profile of adhesion genes, resulting in genetic coupling between the two traits (Fig. S4A). Strictly unicellular and high-adhesion multicellular lineages also often co-regulate cell state and adhesion genes (Fig. S4B), but changes to the expression of adhesion proteins in these cases do not change the adhesion strength between cell states (Fig. 3A, and lineages I, II, IV and V in Fig. S4C).

**Fig. 3.**
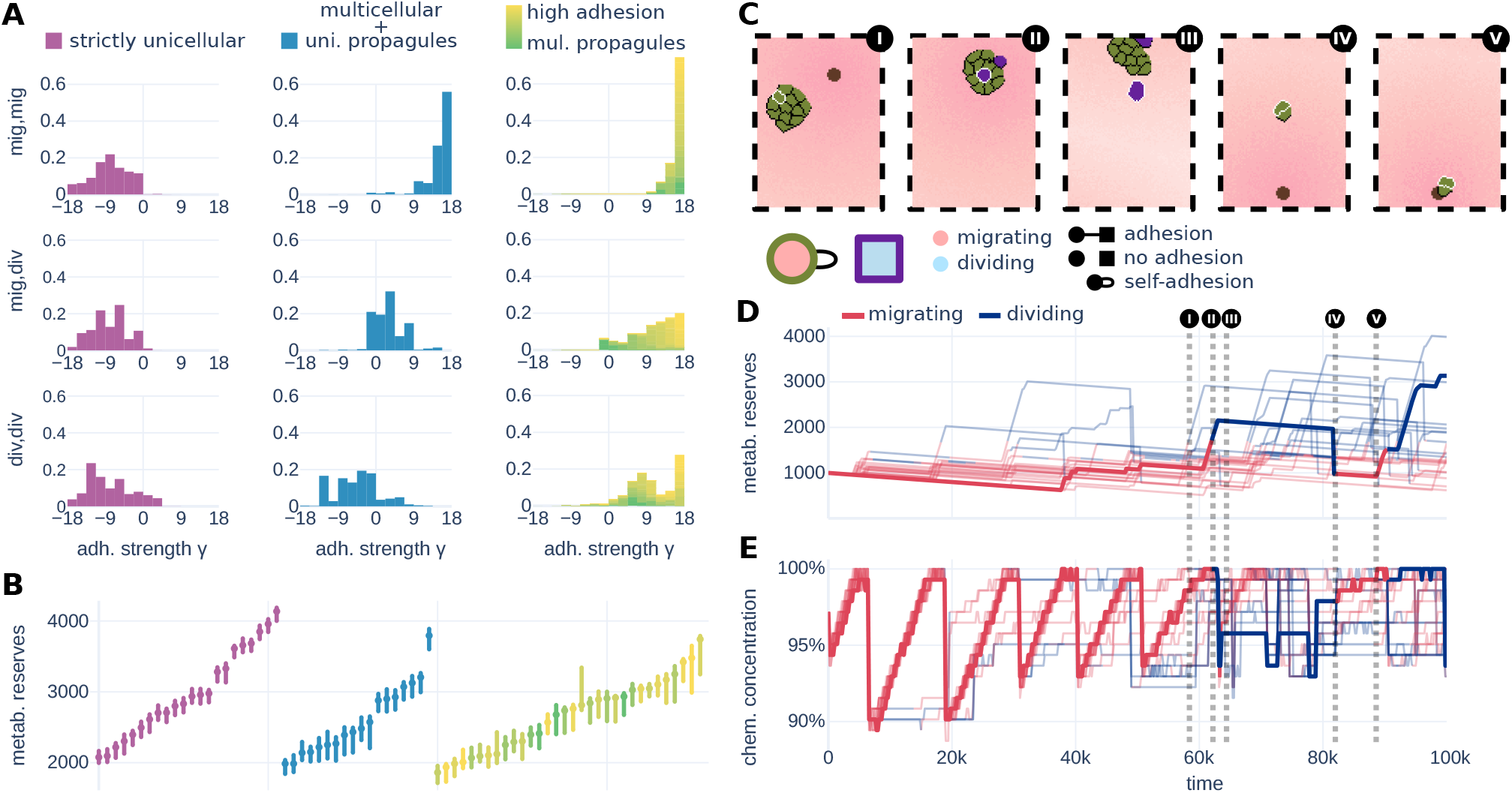
Coordinated expression of cell state and adhesion genes leads to the formation of propagules. **A** Distribution of adhesion strengths (*γ*) measured across our simulations for cells expressing different pairings of cell states. Propagule-forming life cycles couple the regulation of cell state and adhesion proteins, leading to differential adhesion between pairs of cells according to their cell states (note: 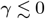 results in cells detaching from one another). **B** Metabolic reserves are used as a cue for division. We show the interquartile range of threshold values beyond which division is initiated for different lineages. Circles mark the medians. **C** Formation of a unicellular propagule. The colour of the cells indicates which combination of adhesion proteins they express. Below the sequence of images, we show a diagram representing how strongly cells expressing each combination of adhesion proteins adhere to each other. The colour inside the symbol shows whether the combination of adhesion proteins is expressed by cells during migration or division. A line connecting the symbols is drawn if cells expressing the two adhesion profiles adhere. Tracking **D** metabolic reserves, and **E** chemoattractant concentration in a subset of cells over time shows how the GRN uses these values to effect adhesion and behaviour changes. The thicker line represents the trajectory of the cell highlighted in white in C.

We next clarify how the observed adhesion dynamics are regulated by the inputs received by the GRN. In all lineages, cells initiate division when their metabolic reserves exceed a specific value. Because this threshold is genetically encoded (Fig. S4), the timing of initiation of cell division varies greatly among lineages, but is very consistent within the cells belonging to each lineage (Fig. 3B). In Fig. 3C, we track a cell within a migrating cluster to show how food consumption activates the coordinated regulation of cell division and adhesion, leading to its transition into a propagule. The tracked cell (white border) initially adheres to its migrating cluster using one adhesion profile (step I, green). After feeding, the cell initiates division and expresses a non-adhering combination of adhesion proteins (II, purple), causing it to sort out from the cluster due to differential adhesion (46) and to detach as a propagule (III). After division, the two daughter cells revert to the original adhesion profile and migrate as a new two-celled cluster (IV) that colonises a new food patch (V). We show that the primary signal used for initiating propagule formation is the level of internal metabolic reserves, which cells use as a threshold to decide when to divide or migrate (Fig. 3D). The chemoattractant signal is used to further tune decision making, causing division to end early when at high concentrations (Fig. 3E, detailed in Fig. S5).

### Widespread co-option drives the evolution of propagules

To investigate the evolutionary steps that lead to propagule formation, we extracted strictly unicellular populations from EC 1 and high-adhesion multicellular populations from EC 6 and let them evolve for an additional 10^8^ time steps in EC 3, which selects for propagule formation according to our previous evolutionary experiments (Fig. 2). This allowed us to analyse the evolution of life cycles under controlled starting genetic landscapes, where all cells are initially unicellular or multicellular. We observed an increase in the adhesion strength between migrating cells 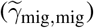 in lineages that were initially strictly unicellular (Fig. 4A, lineage 1), reflecting the establishment of a multicellular migrating phase. Since the adhesion strength between dividing cells and all other cells 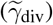 remains low, the lineage is now able to form propagules. Conversely, lineages that were initially high-adhesion multicellular evolve lower 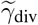 while maintaining high 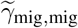, indicating that they too have evolved propagule formation (Fig. 4A, lineage 2).

**Fig. 4.**
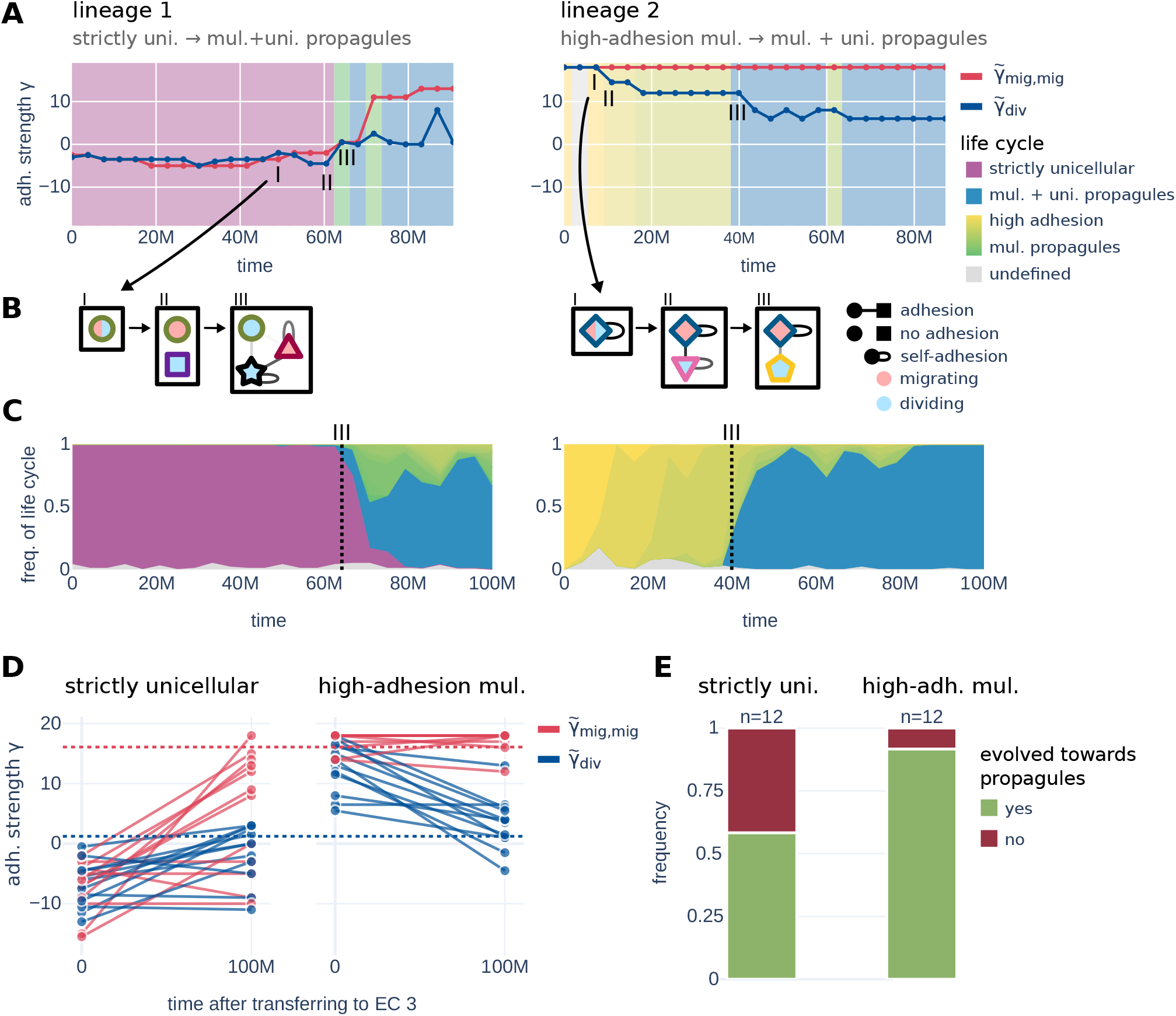
Propagule formation evolves through the co-option of unicellular or multicellular life cycles. **A** Evolution of adhesion properties of a strictly unicellular lineage taken from EC 1 (lineage 1) and a strictly multicellular lineage taken from EC 6 (lineage 2) after their cells are transferred to an environment selecting for propagule formation (EC for 10_8_ time steps. The median adhesion strength between migrating cells 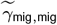 and between dividing cells and all other cells 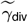 is shown in red and blue, respectively, with the background colour indicating the life cycle expressed by the lineage at that time (see Materials and methods, Categorisation of life cycles). **B** Evolution of regulation of adhesion proteins. Each unique combination of adhesion genes expressed by a lineage is represented by a unique colour and symbol (same as Fig. 3C). The inner colour of the symbols indicates whether that combination of adhesion genes is expressed during division or migration. A line connecting the symbols is drawn if cells expressing the two adhesion profiles adhere. **C** Distribution of life cycles among 100 cells taken from the simulations where lineages 1 and 2 originated. Time step III marks the first time propagule formation evolved in lineages 1 and 2. **D** Evolution of 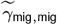 and 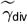 of strictly unicellular lineages (left) and high-adhesion multicellular lineages (right) after they were transferred to EC 3. Dotted lines indicate the average value for the trait in populations endogenous to EC 3. **E** 7 out of the 12 lineages that started with a strictly unicellular life cycle (left) and 11 out of the 12 lineages that started with a high-adhesion life cycle (right) evolved towards making propagules (respectively by increasing their 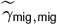 and decreasing their 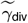, see subfigure D) after being transferred to EC 3.

Following individual lineages reveals that propagule formation evolves via co-option of the lineage’s ancestral life cycle. Cells in lineage 1 (Fig. 4B) initially display a strictly unicellular life cycle after being transferred from EC 1 to EC 3, with cells expressing a single adhesion profile (I). The first step towards forming propagules consists of evolving genetic coupling between cell state and adhesion proteins (II): the ancestral adhesion profile (green circle) is now expressed only during migration, while a new profile (purple square) is expressed during division. This change has no immediate functional effect, since the new adhesion profile does not result in intercellular adhesion. Nevertheless, it represents a pre-adaptation for the subsequent evolution of propagule formation. The critical step to the evolution of propagules (III) is the appearance of adhesion proteins that cause migrating cells to adhere to each other (dark red triangle), while the expression of the ancestral (green circle) adhesion proteins becomes associated with the dividing cell state. Because there is no adhesion between cells expressing these two adhesion profiles, dividing cells can separate from migrating clusters to form propagules. Once evolved, propagule formation takes over the unicellular population and fixes rapidly (Fig. 4C), highlighting its selective advantage in EC 3. In lineage 2, propagules also evolve via co-option, this time of an ancestral multicellular life cycle. The ancestral combination of adhesion proteins (dark blue diamond) is co-opted for the regulation of a migrating phase, followed by the evolution of novel combinations of adhesion proteins (pink triangle and yellow pentagon) that enable propagule formation (4B). The evolution of this trait in lineage 2 also leads to a fast selective sweep in the population (Fig. 4C). In Fig. S6, we show that unicellular propagules evolved via co-option in 5 out of 7 lineages that were originally strictly unicellular, and in 7 out of 8 lineages that were originally high-adhesion multicel-lular, according to our detection algorithm (see SI Appendix, Inference of co-option).

Strictly multicellular lineages seem to have evolved propagule formation more easily than strictly unicellular lineages when transferred to EC 3, evidenced by the fact that 11/12 multicellular lineages evolved a lower 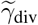, while only 7/12 strictly unicellular lineages evolved a positive 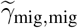 in the same period of time (Fig. 4D and E). Presumably, this disparity in the number of lineages that evolve propagules is due to the difference in mutation availability between the two cases. Starting from a non-adhering state, most regulatory mutations do not lead to changes in adhesion, creating a large neutral plateau in phenotype space. Thus, strictly unicellular lineages may experience a long series of neutral mutations before reaching a combination that produces adhesion, at which point mutations affecting adhesion proteins become selective. Altogether, this suggests that the initial evolution of adhesion reshapes the accessible evolutionary pathways, facilitating the evolution of subsequent multicellular traits such as propagule formation.

### Eco-evolutionary protection of multicellularity

To better understand how environment colonisation and competition could shape ecological dynamics at the onset of multicellularity, we ran a series of experiments where lineages with different life cycles invade ECs 1, 3 and 6 and compete with host lineages that evolved in these environments. (Fig. 5A). In most instances, the host lineages — which evolved in the environment where competition is taking place — out-competed the invading lineages. However, invading lineages with a multicellular + unicellular propagule life cycle out-competed host lineages in EC 1 in all replicas of the experiment. This prompted us to investigate whether multicellularity could represent a viable strategy in EC 1, despite not evolving there (Fig. 2).

**Fig. 5.**
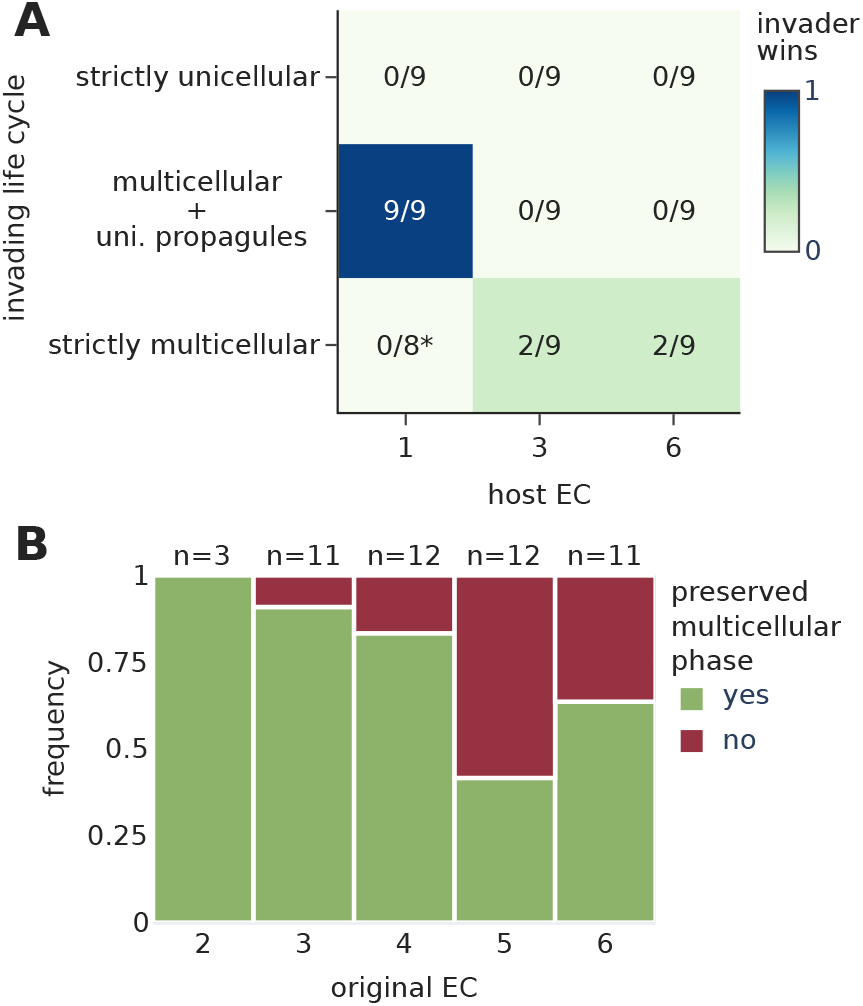
Multicellularity is evolutionarily protected in EC 1. **A** Number of competitions where the invading strategy wins against the host strategy. We compete three invading lineages with different life cycles against three host lineages in ECs 1, 3, and 6. Lineages were selected based on their inferred fitness (see Materials and methods, Inference of fitness) in the environment hosting the competition. The asterisk marks one data point that we excluded because neither lineage outcompeted the other in the simulation. Mutations were disabled in these simulations to prevent further evolution. **B** Frequency of initially multicellular strategies that preserved a multicellular phase after being transferred to EC 1 for 10_8_ time steps.

After transferring strictly multicellular and multicellular + unicellular propagule lineages to EC 1 and letting these lineages evolve for 10^8^ time steps, we observed that multicellularity persisted in the majority of the simulations (35/49, Fig. 5B), with all lineages either preserving or evolving a propagule-forming life cycle (Fig. S7A). Moreover, multicellularity was better protected in lineages that had originally evolved in environments similar to EC 1, as evidenced by the large number of lineages from ECs 5 and 6 that lost multicellularity after the transference. This suggests that protection of multicellularity may be contingent on certain phenotypic or genetic factors. For example, reaching the basin of attraction of the multicellular life cycles that perform well in EC 1 might be easier if the lineage can already make propagules or is pre-adapted to environments with a fast turnover of food patches.

In the symmetric experiment, where we transfer strictly unicellular and multicellular + unicellular propagule lineages to EC 6, the inverse pattern can be observed, with a unicellular phase being preserved more often in the ECs that are most different from EC 6 (ECs 1 and 2 in Fig. S7B and C), rather than similar. After being transferred to EC 6, the lineages that preserved a unicellular phase displayed poor fitness. This is also unlike the previous experiment, where the lineages that preserved a multicellular phase performed well in their new environment (compare Fig. S7D and E). Altogether, this suggests that lineages preserving a unicellular phase did not adapt well to EC 6, most likely because they struggled to navigate the phenotype space due to low mutational availability (similarly to the case shown in Fig. 4D and E).

## Discussion

Understanding the origin of multicellular reproduction requires explaining how traits of individual cells were repurposed to organise development (43, 44). Here, we show that developmental regulation arises spontaneously through adhesion-mediated interactions between neighbouring cells. In our computational framework, cells can evolve a gene regulatory network to interact with other cells and the environment, but we do not impose developmental rules or ecological responses as a selection criterion. This allows us to investigate the evolution of reproduction as a novel, emergent process (57). We find that both unicellular and multicellular modes of reproduction could evolve in our simulations, depending on the heterogeneity of the distribution of food in the environment. In intermediate environments, cell lineages evolved multicellular clusters that periodically release propagules. Propagules are beneficial in these environments because they improve dispersal, while the cluster from which they originate benefits from collective chemotaxis. We show that these lineages regulate the formation of propagules by coupling their cell state to specific expression profiles of adhesion proteins. This trait evolves through the co-option of ancestral strictly unicellular or strictly multicellular life cycles, which are encapsulated as a phase in the developmental cycle of the propagule-forming strategy.

Comparative genomic and phylogenetic studies have shown that the evolution of several multicellular traits involved the co-option of proteins that originally served different functions in their unicellular ancestors (37, 53, 58). In our simulations, multicellular reproduction evolved through the same process. The architecture of the GRN is fixed in our model, and no new regulatory machinery can evolve. Yet, multicellular reproduction still emerged via the co-option of modules of regulation of adhesion. Before the evolution of multicellularity, these modules mediated ecological interactions between single-celled organisms, which spread apart in the environment by expressing adhesion proteins with low complementarity. The same regulatory modules were then repurposed during the transition towards multicellularity to regulate the formation of propagules used for reproduction in a developmental context. Our model therefore highlights co-option as a mechanism through which properties of individual cells can be integrated into coordinated reproductive cycles featuring a unicellular propagule stage. These propagule-forming life cycles bear a striking similarity to those observed in previous experiments performed in *C. reinhardtii*, where the rapid evolution of propagules was suggested to be evidence of co-option (17). Our results corroborate this hypothesis, showing that co-option of ancestral life cycles for reproduction is not only possible, but evolutionarily likely. More broadly, our results suggest that co-option might represent a general solution to the emergence of collective traits, enabling the evolution of complex life cycles while avoiding the organisational costs associated with building more elaborate regulatory machinery.

The eco-evolutionary feedback between cells and sources of food in our model recapitulates real-life examples of ecology shaping the evolution of life cycles, and therefore provides a possible explanation for their evolution. Organisms that alternate between unicellular and multicellular lifestyles often rely on fluctuations of nutrient availability to regulate switches between these stages. Similarly to how propagule-forming lineages in our model migrate as a multicellular cluster during a feeding stage and then produce propagules after food is depleted, many amoebozoans and opisthokonts perform collective feeding when resources are abundant, followed by unicellular reproduction through single-celled structures once food becomes scarce (14). For example, the holomycotan *Fonticula alba* forms cell collectives that feed on patches of bacteria and later differentiate into a sorocarp for reproduction through spores (30). Collective feeding fol-lowed by unicellular reproduction has also been described in Myxogastria (59–61), Labyrinthulomycota (62, 63), and choanoflagellates like *Salpingoeca rosetta* (64–66). Other choanoflagellates, e.g. *Choanoeca flexa*, instead use alternative environmental signals such as salinity to regulate a transition to the multicellular stage (67). Interestingly, this species forms multicellular groups both through aggregation and continued cell division, similarly to the cells in our simulations.

We showed that multicellularity could not initially evolve in environments with homogeneous resources (EC 1) when starting from random genomes. This may be due to unicellular strategies being more adaptive than early forms of multicellularity in these environments. Once lineages become unicellular, mutations that lead to intercellular adhesion become rare, making multicellularity less likely to evolve (Fig. 4D and E). However, this ecological interference between unicellularity and weak multicellularity could be avoided by allowing lineages to first evolve multicellularity and propagules in intermediate environments (EC 3), after which they could successfully colonise EC 1. Similar effects have been previously described in the context of protein evolution, where changes in the environment can bridge fitness valleys and enable further functional adaptation of a protein sequence (68–70). Our results suggest that environmental shifts can likewise promote the evolution of major innovations, such as multicellularity. The role of EC 3 in facilitating this transition is reminiscent of environmental scaffolding, where particular ecological contexts transiently support the emergence of new levels of organisation (71–75).

In summary, we have shown that variation in the spatial distribution of resources in the environment promotes the evolution of diverse life cycles and modes of reproduction. Our model captures this constructive form of evolutionary innovation (57), showcasing how ecological interactions can be co-opted to regulate development at the onset of multicellularity.

## Materials and methods

Our model consists of an evolving population of cells living in a two-dimensional arena. Cells must consume resources to survive and replicate. Resources are patchily distributed and release a chemical signal that cells can follow through chemotaxis. Each cell contains a GRN represented as a Boolean network. The expression of different genes in the GRN determines whether the cell is migrating or dividing and whether it adheres to neighbouring cells.

Following the CPM formalism, we implement a Hamiltonian equation which describes the energy of the system. A Monte Carlo method is then used to minimise this energy by accepting or rejecting copy attempts between pairs of lattice sites. A full description of the model can be found in the SI Appendix, Model description. The source code of the model was built on top of the Tissue Simulation Toolkit framework (45) and can be found at https://github.com/aleferna12/regulation_evolution2. Parameters used for the simulations are available in Table S2.

### Non-evolutionary simulations

After obtaining the results from our evolutionary simulations (as described in Results), we ran a set of non-evolutionary simulations where mutations are disabled. The purpose of these simulations was to build a large, fine-grained dataset that we could use to analyse functional traits of the life cycles that evolve in our model. We start each simulation with 100 cells cloned from an evolved lineage’s most recent common ancestor (MRCA). The simulation then runs for 10^6^ time steps while the cell population reaches steady state, after which we record the data used for the analyses described below (over the course of an additional 10^6^ time steps). Data points are recorded every 20,000 time steps.

### Categorisation of life cycles

We categorise the life cycles evolved in the simulations into three groups: strictly unicellular, multicellular + unicellular propagules, and strictly multicellular. To classify a lineage’s life cycle, we first record all unique combinations of adhesion proteins expressed by its cells in a non-evolutionary simulation (see Materials and methods, Non-evolutionary simulations), storing them in the vector *E*. We also store the frequency with which each combination of adhesion genes was expressed by the cells according to their cell state in the vectors *F* ^mig^ and *F* ^div^, such that 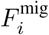 is the frequency with which a cell in the simulation is both migrating and expressing *E*_*i*_. We then calculate the median adhesion strength between cells, given their cell states:

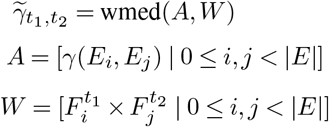

where *t*_1_ and *t*_2_ are cell states (either migrating, dividing, or all, in which case *F* ^all^ =*F* ^mig^ +*F* ^div^), wmed(*A, W* )gives the median of the vector of adhesion strengths *A* weighted by the vector of expression frequencies *W*, and *γ*(*E*_*i*_, *E*_*j*_)is the adhesion strength calculated by comparing the vectors of expression of adhesion proteins *E*_*i*_ and *E*_*j*_ expressed by cells with sigmas *s*_*i*_ and *s*_*j*_ respectively, such that *γ*(*E*_*i*_, *E*_*j*_)=*γ*(*s*_*i*_, *s*_*j*_)(see SI Appendix, Cell adhesion).

This gives us six combinations of adhesion strength 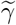 values between the different cell states: 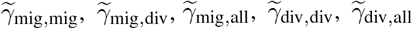, and 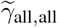 We usually abbreviate 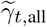 to simply 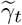, for example 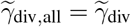 in the text and figures. We then classify a life cycle according to the following decision tree: if migrating cells do not adhere to other migrating cells (i.e.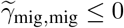), we classify the life cycle as strictly unicellular. Else, if dividing cells do not adhere to other dividing cells (i.e.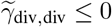), we classify the life cycle as multicellular + uni cellular propagules. Otherwise, we classify the life cycle as strictly multicellular, and use 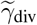 to differentiate between lineages that often form propagules (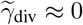, green in Fig. 2) and those that never or rarely do (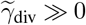, yellow in Fig. 2).

In Fig. 4, we examine the evolution of propagule formation across 10^8^ time steps. Because the evolutionary simulation contains only a few cells of the lineage of interest, we cannot directly extract enough data to run the analysis as described above. Instead, to calculate the *γ* values displayed in Fig. 4, we artificially feed the GRN of the lineage’s ancestor with inputs experienced by other cells in the simulation, and compare the GRN’s outputs with data from the population in real-time during evolution. When a life cycle cannot be reliably identified with this method, we conservatively label it “undetermined” in the figure.

### Measure of dispersal

To measure the dispersal of a lineage in its environment, we periodically record the positions of all cells in the lattice in a non-evolutionary simulation (see Materials and methods, Non-evolutionary simulations). We then perform a standard quadrat analysis using a quadrat size of 100×100 lattice sites. We define the dispersal of the population at a time step *t* as:

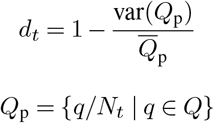

where 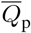 is the mean of the set containing the proportion of cells in each quadrat *Q*_p_, *N*_*t*_ is the population size at time *t*, and *Q* is a set containing the number of cells on each quadrat at time step *t*. This ensures that our dispersal metric *d*_*t*_ lies in the interval (0, 1], where *≈* 0 represents the situation where all cells are contained in a single quadrat and 1 represents the situation where each quadrat contains the same number of cells. Finally, we define the dispersal of the lineage as the mean of all of its recorded *d*_*t*_ (measured every 20,000 time steps in non-evolutionary simulations, see Materials and methods, Non-evolutionary simulations).

### Inference of fitness

We infer the fitness of a population by measuring how much food it has consumed from its environment at steady state (when the rate of food consumption and the influx rate of food reach equilibrium). Populations that can empty the environment of food before reaching steady state are considered to have fitness 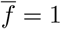, while those that require the lattice to be full to survive have fitness 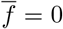. 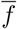 is calculated by taking the mean of the time series of *f*_*t*_, measured every 20,000 time steps in non-evolutionary simulations (see Materials and methods, Non-evolutionary simulations), where *f*_*t*_ =1*− β*_*t*_/Γ, with *β*_*t*_ being the number of food units on Λ_food_ at time step *t* and Γbeing the maximum amount of food supported on the lattice.

## ACKNOWLEDGEMENTS

This work was supported by grants from the Gatsby Charitable Foundation (G112566) to R.M.A.V. and from the Cambridge Trust to A.P.F. We thank members of the Sainsbury Laboratory for helpful discussions and suggestions.

