## supplemental information for "Reproduction emerges from ecological interactions at the onset of multicellularity"

March 9, 2026

### Model description

Cells are modeled in a square lattice $\Lambda_{\text{cell}} \subset \mathbb{Z}$ of size $L^2 = 2000 \times 2000$ following the Cellular Potts Model formalism (1, 2). A cell $c$ is composed of lattice sites $x$ that share the same spin $s$, i.e. $c(s) = \{x \mid x \in \Lambda_{\text{cell}} \wedge \sigma(x) = s\}$. The configuration of spins on the lattice is updated stochastically according to a Metropolis algorithm. Each time step (used interchangeably with Monte Carlo step), we randomly sample $L^2$ sites $x$ from the lattice and select a site $x' \in$ neigh(x) in their Moore neighborhood, which may copy its spin $\sigma(x')$ into $x$. The probability that this copy attempt is successful depends on the difference that it would incur to the energy $H$ of the system, described by a Hamiltonian function, as well as the migration biases $Y(x, x')$:

$$H = H_{\text{size}} + H_{\text{adh}}$$

$$Y(x, x') = Y_{\text{pers}}(x, x') + Y_{\text{chem}}(x, x')$$

If a copy attempt would result in a decrease in the total energy of the system, i.e. $\Delta H + Y(x, x') < 0$, it is always accepted. Otherwise, a Boltzmann distribution is used to calculate the probability that the copy attempt might still succeed due to random "thermal" fluctuations in the environment:

$$P\left(\Delta H, Y(x, x')\right) = e^{-\frac{\Delta H + Y(x,x')}{T}}$$

where $T = 16$ is the Boltzmann temperature. To increase computational efficiency, we restrict sampling to cell boundaries following the implementation of van Steijin et al. (3).

#### Cell size maintenance

The area of a cell $A(c) = |c|$ fluctuates around the target area parameter $A_{\text{t}} = 50$. Deviations from this target increase the system's total energy (represented by the term $H_{\text{size}}$ in the Hamiltonian equation):

$$H_{\text{size}} = \sum_{c \in C} \lambda \times (A(c) - A_{\text{t}})^2$$

where $C$ is the set of cells and $\lambda = 4$ is the resistence of cells to deviations in area.

#### Cell adhesion

The total contact energy resulting from interactions between neighboring cells is implemented as the $H_{\text{adh}}$ term in the Hamiltonian:

$$H_{\text{adh}} = \sum_{(x,x')} J(\sigma(x), \sigma(x')) \times (1 - \delta(\sigma(x), \sigma(x')))$$

$$\delta(a,b) = \begin{cases} 1 & \text{if } a = b \\ 0 & \text{otherwise} \end{cases}$$

where $(x, x')$ are all pairs of neighboring sites on the lattice, such that $x' \in$ neigh(x). Calculations are restricted to cell boundaries by the Kronecker delta $\delta$. The contact energy of a pair of sites with different spins $J(s_1, s_2)$ is proportional to the similarity of the vectors of expression states of ligand $I(s)$ and receptor $R(s)$ genes of the cells associated with the two spins:

$$J(s_1, s_2) = J_\alpha + \mathrm{sim}(R(s_1), I(s_2)) + \mathrm{sim}(R(s_2), I(s_1))$$

$$\mathrm{sim}(R, I) = \sum_{i=1}^{\nu} i\delta(R_i, I_i)$$

where $J_\alpha = 12$ is the minimum contact energy between any two cells and $\nu$ is the number of receptor and ligand genes of the cells, and $R_i$ and $I_i$ are the expression states (0 or 1) of the $i^{\text{th}}$ receptor and $i^{\text{th}}$ ligand, respectively. We assume that the medium is inert and that the contact energy between a cell and the medium is constant, such that in the special case where either $s_1$ or $s_2$ is the spin that represents the medium $s_{\text{med}}$, then $J(s, s_{\text{med}}) = J'_\alpha$, where $J'_\alpha$ is a parameter with value 24. Thus, the strength $\gamma(s_1, s_2)$ with which two cells with spin $s_1$ and $s_2$ adhere can be described in terms of the difference between the contact energy of cells and the medium and the contact energy between the two cells:

$$\gamma(s_1, s_2) = J'_\alpha - \frac{J(s_1, s_2)}{2}$$

For our simulations, we chose values $J_\alpha$, $J'_\alpha$, and $\nu$ such that the range of possible $\gamma(s_1, s_2)$ between two cells is $[-18, 18]$, with negative values indicating that the cells do not adhere, and positive that they do.

#### Food and metabolism

Food is modeled in a separate lattice $\Lambda_{\text{food}}$ with the same dimensions as $\Lambda_{\text{cell}}$. Throughout the simulation, we introduce food in $\Lambda_{\text{food}}$ every $1/\phi$ time steps as a "patch" with area $A_{\text{food}}$ in the approximate shape of a circle (Fig. 1A). When created, a food patch is composed of 64 layers of food units, representing a multilayer bacterial biofilm. We vary $\phi$ and $A_{\text{food}}$ in our simulations to represent different environmental conditions (see Table 1 and Fig. S1). To decide where a new food patch is created, we query random positions on the lattice until we find one that is no closer than $d_{\text{min}}$ to any food patch currently in the environment:

$$d_{\text{min}} = \frac{L}{\sqrt{2|\mathcal{P}|}}$$

where $L$ is the size of the lattice and $\mathcal{P}$ is the set of food patches currently in the environment. We then center the new food patch at this position. This is done to ensure that food patches are never created too close to each other and that cells always need to migrate across the environment to reach them. We set a maximum number of food units $\Gamma$ allowed in $\Lambda_{\text{food}} = 524288$, such that a food patch creation attempt is canceled if it would result in exceeding this limit. This is done for computational reasons, as the algorithm that selects a semi-random position for the food patch described above has a high rejection ratio when the food patch density is very high.

Cells that share a site $x$ on $\Lambda_{\text{cell}}$ with a corresponding site containing food $y$ on $\Lambda_{\text{food}}$ can consume one food unit from $y$ provided that they have not eaten in the last $\tau_{\text{eat}} = 2$ time steps. When consumed, the food unit is removed from the lattice and added to the cell's metabolic reserves. Every $\tau_{\text{met}} = 100$ time steps, we decrease the metabolic reserves of all cells by one unit of food, representing a metabolism. Therefore, the ratio $\frac{\tau_{\text{met}}}{\tau_{\text{eat}}}$ determines the rate at which cells can accumulate metabolic reserves. Cells that run out of metabolic reserves are considered dead and are removed from the lattice. Additionally, all cells have a probability $\zeta = 5 \times 10^{-6}$ of dying each time step regardless of how much food they have accumulated. This probability accounts for death caused by processes we do not model explicitly, such as metabolic stress.

#### Gene regulation

Each cell in a simulation contains a GRN with a fixed topology composed of three layers of genes that control its behavior (Fig. 1B). The first layer consists of two sensory genes that provide the cell with information about (i) the local concentration of the chemoattractant signal that emanates from the food sources and (ii) the amount of metabolic reserves the cell has left to spend. These sensory genes are connected to a middle layer of $2\nu + 2 = 18$ regulatory genes representing a network of transcription factors that control each other's expression as well as the expression of the functional genes. Lastly, an output layer contains the functional genes that determine cell behavior. The first of these genes determines the cell state (whether the cell is currently migrating or dividing), followed by $\nu$ ligand genes and $\nu$ receptor genes that together determine how strongly the cell adheres to its neighbors.

The expression state of regulatory and output genes in the network is boolean (either 0 - not expressed or 1 - expressed). Regulatory genes receive input both from the sensory genes (i.e. the chemical gradient and metabolic reserves of the cell) and from all regulatory genes, including themselves. Functional genes receive input only from genes in the regulatory layer. Every $g = 20$ time steps, chosen so we even out microscopic fluctuations in the cell parameters, we synchronously update the expression state of each gene $i$ in the network as follows:

$$r_i(t+1) = \sum_{j \in \mathcal{N}_i^-} w_{j \to i} S_j(t)$$

$$S_i(t+1) = \begin{cases} 1 & \text{if } r_i > \rho_i \\ 0 & \text{otherwise} \end{cases}$$

where $\mathcal{N}_i^-$ is the set of in-neighbours of $i$ and $r_i(t+1)$ is the total input into gene $i$ coming from genes $j$ with state $S_j$ at time $t$, which respectively regulate $i$ with weight $w_{j \to i}$. $S_i(t+1)$ is then the new state of gene $i$, determined by whether $r_i$ is larger than $\rho_i$, the threshold of activation for this gene.

#### Division

A cell will start division if the state of the first functional gene of its GRN is 1 for twenty consecutive GRN updates (corresponding to $\eta_{\text{init}} = 400$ time steps), representing a period during which the cell prepares its cellular machinery before the beginning of division. Once division starts, the cell can no longer migrate or perform chemotaxis, simulating the conflicting use of the cytoskeleton for division and migration characteristic of some microorganisms (4, 5). During division, we progressively increase the target area of the cell at a fixed rate for $\eta_{\text{grow}} = 2 \times 10^4$ time steps to $2A_{\text{t}}$, causing the cell to grow approximately twice its original size. At the end of this period, we divide it along its minor axis to form two daughter cells. Division is halted if the cell expresses 0 in its cell state gene at any point during this process, which will reset its cell state to migrating. When the cell divides, each weight and activation threshold in the GRN of one of the daughter cells has a $m = 0.05$ probability of mutating. If a mutation event occurs, we then add a small random number sampled from a normal distribution with mean 0 and $\sigma_{\text{mut}} = 0.05$ to the mutated network parameter. The other daughter cell inherits its genome from the mother unaltered. The metabolic reserves accumulated by the mother cell are equally partitioned between the daughters.

#### Migration

When the state of the cell state gene is 0 (or if it has been 1 for less than $\eta_{\text{init}}$ time steps), the cell is in migrating mode. We model persistent migration by biassing the movement of a cell with spin $s$ towards its previous direction of motion $\psi(s)$:

$$Y_{\text{pers}}(x, x') = -\mu_{\text{pers}} \cos(\psi(\sigma(x')) - \theta(s, x))$$

where $\theta(s, x)$ is the angle between the center of mass of the cell with spin $s$ and the target site of the copy attempt, and $\mu_{\text{pers}}$ is the maximum energy bias of persistent migration, which we set to 3 for migrating cells and to 0 for dividing cells, making the latter stationery. $\psi(s)$ is updated every 50 time steps with the actual direction of motion of the cell during this period, resulting in cells that move consistently in the same direction over short time scales and display Brownian motion over long scales (6).

#### Chemotaxis

Migrating cells can also perform chemotaxis by following a noisy chemical signal that emanates from the sources of food, forming a gradient that decays linearly. We do not model diffusion for computational efficiency, and instead assume that the gradient is static. We model this chemical signal in a separate lattice $\Lambda_{\text{chem}}$ with the same dimensions as $\Lambda_{\text{cell}}$. The concentration of the chemical signal $\chi(x)$ at site $x$ is determined by its distance to the closest food patch:

$$\chi(x) = 1 + k_\chi \times (D - d_{\text{food}}(x))$$

$$d_{\text{food}}(x) = \min(\{d(x,p) \mid p \in \mathcal{P}\})$$

where $k_\chi = 0.05$ is a scaling constant which determines the slope of the gradient, $D$ is the diagonal of the lattice, and $d(x,p)$ is the Euclidean distance between $x$ and the center of the food patch $p$. To add noise to the chemical gradient, we truncate non-integer values of $\chi$ to their integer part $\lfloor \chi \rfloor$ with probability $\chi - \lfloor \chi \rfloor$. Otherwise, they are changed to the closest integer larger than $\chi$. The contribution of a food patch $p$ to the chemoattractant gradient is removed from $\Lambda_{\text{chem}}$ once the last food particle of $p$ is eaten.

To implement chemotaxis, we find the signal's center of mass by averaging $\chi$ across the cell lattice sites. The chemoattractant direction is then given by the angle $\omega(s)$ spanning from the center of mass of a cell with spin $s$ to the center of mass of the signal perceived by that cell. Chemotaxis occurs because the success rate of lattice site copy attempts performed by a cell is larger if their direction aligns with $\omega(s)$:

$$Y_{\text{chem}}(x, x') = -\mu_{\text{chem}} \cos(\omega(\sigma(x')) - \theta(s, x))$$

where $\mu_{\text{chem}}$ is the maximum energy bias of chemotaxis, which we set to 1 for migrating cells and to 0 for dividing cells. Note that the speed of a single cell's chemotaxis depends on the slope of the gradient (due to averaging X), which is constant across simulations.

### Selective advantages of multicellularity and unicellularity

To confirm that collective migration enhances survival rates in heterogeneous environments, we transfer strictly unicellular lineages to EC 6. In this environmental condition, the inability of strictly unicellular lineages to form multicellular clusters results in slower migration and high death rates due to starvation (Video S5, Fig. S2A and B). Conversely, to investigate the effect of higher dispersal rates on homogeneous environments, we placed high-adhesion multicellular lineages in EC 1, showing that they struggle to take advantage of the food available because the clusters cannot easily divide and spread through the environment, forcing all of the constituent cells to compete for the same resource-poor food patches (Video S6, Fig. S2C and D).

### Inference of co-option

To determine whether unicellular propagules evolved through co-option in lineages that were transferred from EC 1 or 6 to EC 3, we tracked how their adhesion profiles evolved through time. We start by extracting 25 individuals from the lineage's ancestry at regular time intervals of approximately $3.5 \times 10^8$ time steps, recording the adhesion profiles they express through their life cycle. Then, starting from the MRCA, we check for each individual whether the adhesion profile of interest $p$ has a homologous adhesion profile $p_\mathrm{h}$ in the ancestor of that individual, until we find the last individual that displays the ancestral life cycle of the lineage (either strictly unicellular or high-adhesion multicellular). If homology is preserved throughout, including in this individual, then we say co-option occurred.

We choose the initial adhesion profile $p$ in the MCRA differently based on whether the lineage had a strictly unicellular or high-adhesion multicellular life cycle before evolving unicellular propagules. If the lineage was initially unicellular, we pick the adhesion profile most frequently expressed by the propagule stage of the MRCA (e.g. green circle at time step III of lineage 1 in Fig. 4B). Otherwise, we pick the adhesion profile most frequently expressed by the MRCA during collective migration (e.g. blue diamond at time step III of lineage 2 in Fig. 4B). When unicellular propagules evolve from a high-adhesion lineage, it is not always possible to precisely determine when the last individual with the ancestral life cycle lived. This is because the lineage might have at some point evolved multicellular propagules, which cannot be clearly differentiated from high-adhesion multicellular life cycles (Fig. 2A). To circumvent this issue, we conservatively do not halt computation until reaching the last common ancestor of the lineage at time step 0 if the lineage was originally high-adhesion multicellular, requiring that homology is preserved throughout for co-option to be accepted.

Because each individual may express multiple adhesion profiles during its life cycle, we select the homologous adhesion profile $p_\mathrm{h}$ as the adhesion profile with the lowest Hamming distance to $p$. Furthermore, we require that:

1. The Hamming distance between $p_\mathrm{h}$ and $p$ is $\leq 25\%$.

2. $p_\mathrm{h}$ is expressed when given the same combinations of GRN inputs that lead to $p$ being expressed in the descendant.

3. The expression of $p_\mathrm{h}$ results in the same qualitative adhesion behavior as the expression of $p$. If expression of $p$ leads cells to adhere to each other, so must the expression of $p_\mathrm{h}$ (and the opposite is also true).

Criteria 1 and 2 ensure genetic similarity between $p$ and $p_\mathrm{h}$, while criterion 3 ensures that they are also functionally similar. If no adhesion profile in the ancestor satisfies these requirements, we reject co-option for the lineage.

### Supplementary figures

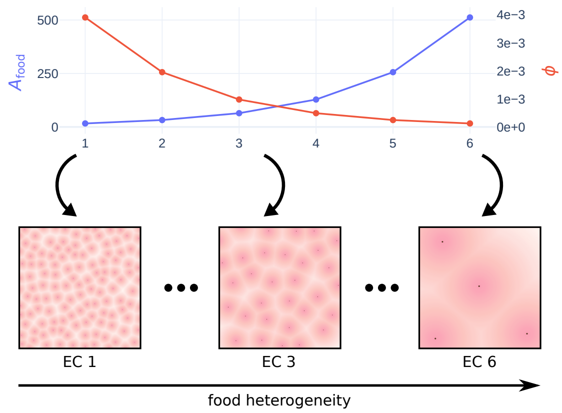


Figure S1: Food heterogeneity as a function of food patch area ($A_{\text{food}}$) and frequency of introduction of food patches ($\phi$) in different Environmental Conditions (EC). For three ECs, we also show a snapshot of the average spatial distribution of food in the environment.

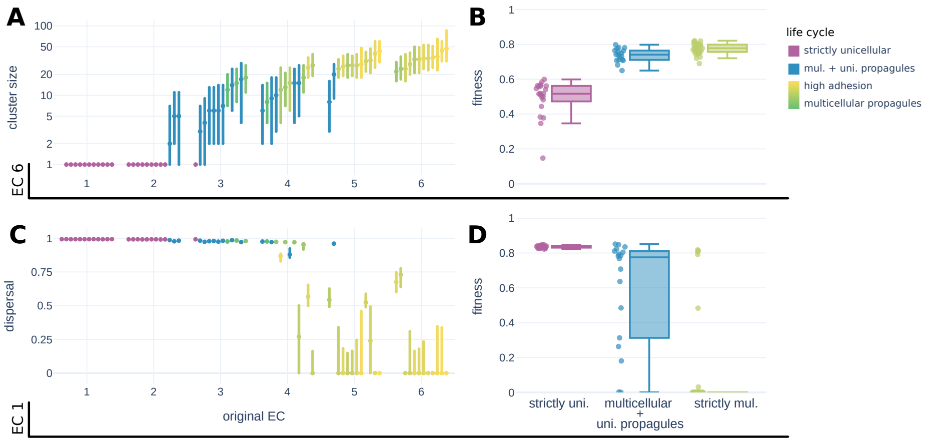


Figure S2: Fitness measurements for lineages after they were transferred across environmental conditions. **A** Interquartile range of cluster sizes and **B** fitness of all lineages after they were transferred to EC 6. **C** Interquartile range of measured dispersal and **D** fitness of all lineages after they were transferred to EC 1. Data was extracted from non-evolutionary simulations (see Materials and methods, Non-evolutionary simulations). See Materials and methods for how dispersal and fitness are measured.

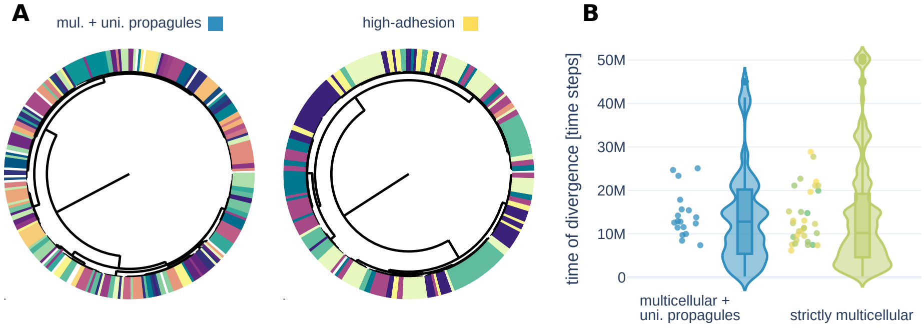


Figure S3: Cluster genetic heterogeneity. **A** Multicellular clusters are composed of many distantly related lineages. We show the phylogeny trees of a representative unicellular propagules lineage (left) and high-adhesion multicellular lineage (right) after evolving $10^8$ time steps in ECs 3 and 6, respectively. Each multicellular cluster in the last time step of the simulations was assigned a unique color, and the leaf nodes of cells belonging to that cluster are painted accordingly. **B** Violin plots of the time since the most recent common ancestor of all pairs of cells that belong to the same cluster in all of our simulations, binned by life cycle. The points represent the means of each lineage when taken independently.

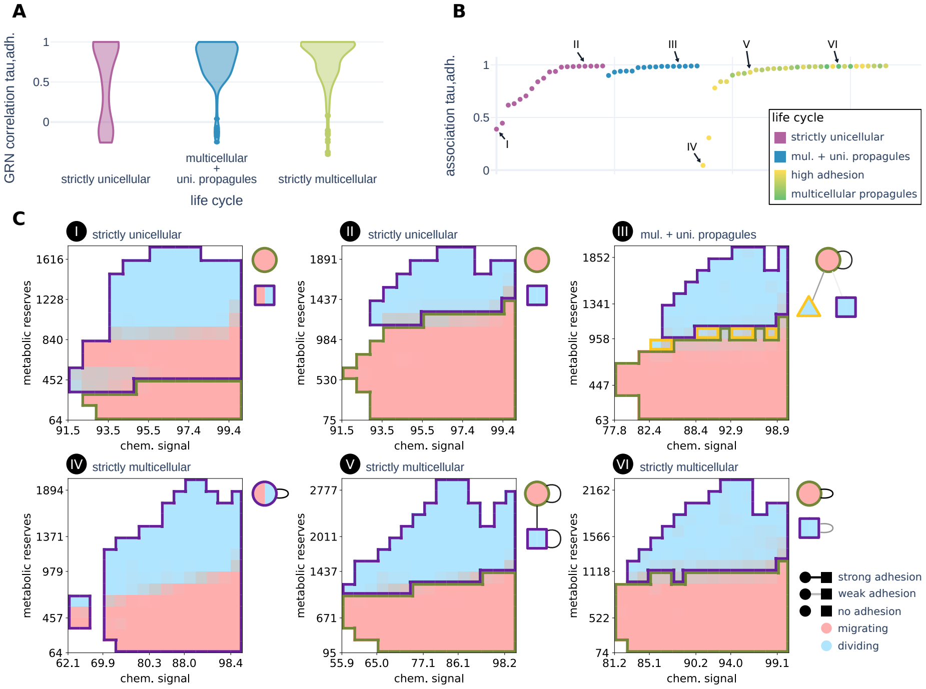


Figure S4: Genetic coupling between cell state (migratory/dividing) and adhesion genes. **A** We measure the influence of a regulatory node $i$ on a functional node $j$ as the conditional probability that $i$ will be expressed given that $j$ is also expressed for all $(i, j)$ pairs of all the evolved lineage's GRNs. We then plot the Pearson correlation between the activation influence of the regulatory nodes on the cell state gene $(j_1)$ and their activation influence on each adhesion gene $(j_2, j_3 \ldots j_{17})$, binning the data points by life cycle. The regulatory genes of strictly unicellular lineages show generally lower correlation between their activation influence on the cell state and adhesion genes, explaining the low association between their expression (subfigure B). **B** Cramer's V association between the expression of the cell state gene and the expression of unique combinations of adhesion proteins (each treated as a different value of a categorical variable). Strictly unicellular and high-adhesion multicellular lineages can have a low association between expression of the cell state and adhesion genes, while this is always high for propagule-forming life cycles. Data was collected from the non-evolutionary simulations (see Materials and methods, Non-evolutionary simulations). **C** Heatmaps showing what cell state and combination of adhesion proteins the cells express in response to different combinations of GRN inputs for a subset of lineages, chosen to reflect the evolved regulatory diversity in different environmental conditions (numbers match those in B). Areas of the heatmaps in red indicate the cells regulate migration for that combination of input parameters, while blue indicates they regulate division. The border colors indicate which unique combination of adhesion proteins the cells express. On the right of each heatmap, we show how strongly cells expressing each combination of adhesion proteins would adhere to each other (similar to Fig. 3). The center of each circle represents whether the combination of adhesion proteins is expressed during migration, division, or both (obtained from the heatmap data).

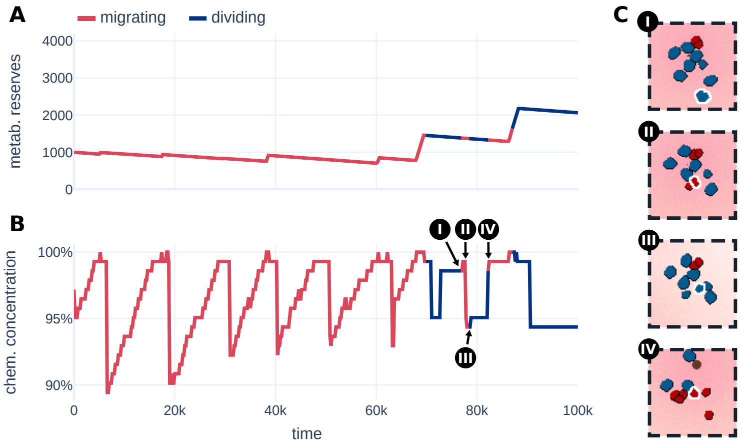


Figure S5: Contribution of the chemoattractant signal for the regulation of the cell state. We show how a cell taken from a simulation that evolved multicellularity + unicellular propagules interprets the **A** metabolic reserves and **B** chemoattractant signal inputs to its GRN to decide when to migrate and divide. Cells can modulate the timing of division in response to the chemotactic concentration: high concentrations can cause early exit from division (II and IV), while low concentrations can lead to early start (III). **C** Frames of the simulation matching the time points indicated in B. The cell highlighted with a white border is the one whose trajectory we follow in A and B.

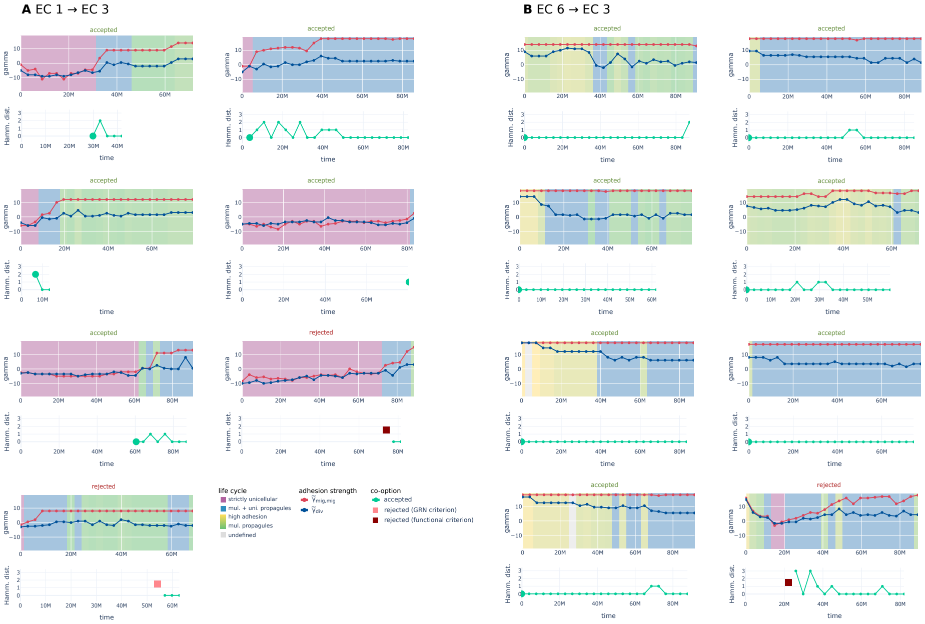


Figure S6: Acceptance and rejection of co-option. We show whether the hypothesis that unicellular propagules evolved via co-option was accepted or rejected for each propagule-forming lineage that evolved in EC 3 after being transferred from **A** EC 1 or **B** EC 6. The evolution of the median adhesion strength between migrating cells $\widetilde{\gamma}_{\mathrm{mig,mig}}$ and between dividing cells and all other cells $\widetilde{\gamma}_{\mathrm{div}}$ is shown in red and blue, respectively, with the background color indicating the life cycle expressed by the lineage at that time. Below, the hamming distance between the ancestral and derived adhesion profile pairs is shown for each iteration of the co-option algorithm (see Inference of co-option). The iteration is terminated either at the last ancestor of the lineage that did not make propagules (represented by a large green circle), in which case co-option is accepted, or if one of the criteria for selecting an ancestral adhesion profile is not met, in which case co-option is rejected. Co-option was rejected for one lineage because there was no overlap in the GRN inputs that produced the ancestral and derived adhesion profiles (pink square, corresponds to criterion 2 in Inference of co-option) and for two lineages because no ancestral adhesion profile was functionally similar to the derived adhesion profile (dark red square, corresponds to criterion 3 in Inference of co-option).

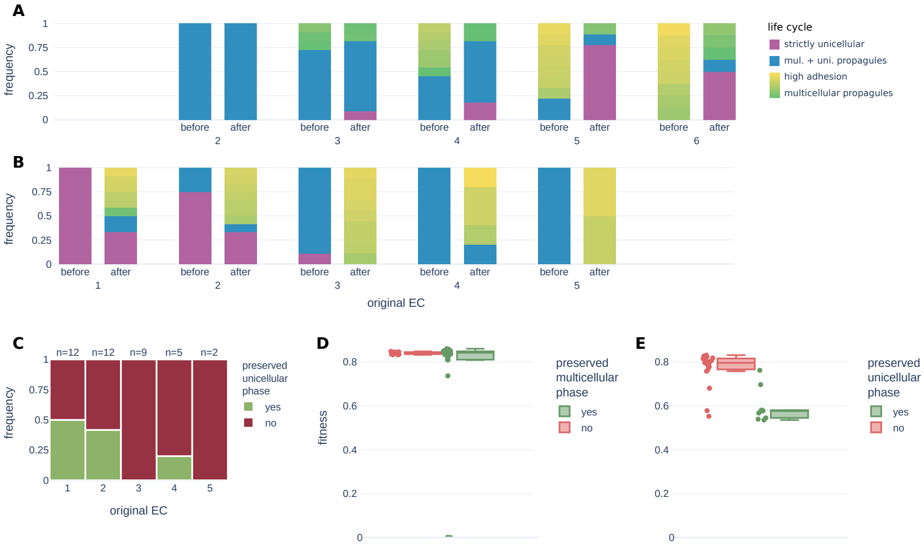


Figure S7: Evolutionary preservation of unicellular and multicellular phases. We show a detailed view of the distribution of life cycles before and after lineages were moved from their original EC to **A** EC 1 and **B** EC 6. **C** Frequency of lineages that were originally strictly unicellular or multicellularity + unicellular propagules and that preserved or lost their unicellular phase after evolving for $10^8$ time steps in EC 6. Fitness change before and after lineages were moved from their original EC to **D** EC 1 and **E** EC 6. Data is binned by whether the lineage preserved a multicellular or unicellular phase, respectively. We only test whether a multicellular phase was preserved for lineages that were not initially strictly unicellular (A and D), and whether a unicellular phase was preserved for lineages that were not originally strictly multicellular (B, C, and E).

### Supplementary tables

| **EC** | $A_{\mathbf{food}}$ | $\phi$ |
|---|---|---|
| EC 1 | $2^4$ | $2^{-8}$ |
| EC 2 | $2^5$ | $2^{-9}$ |
| EC 3 | $2^6$ | $2^{-10}$ |
| EC 4 | $2^7$ | $2^{-11}$ |
| EC 5 | $2^8$ | $2^{-12}$ |
| EC 6 | $2^9$ | $2^{-13}$ |

Table 1: Parameters of the six environmental conditions.

| Parameter | Explanation | Value |
|---|---|---|
| $L$ | size of the square lattice | 2000 lattice sites |
| $T$ | Boltzmann temperature | 16 AUE |
| $\lambda$ | cell stiffness | 4 AUE/[lattice site]$^2$ |
| $A_t$ | target area of a cell | 50 lattice sites |
| **Cell adhesion** | | |
| $J_\alpha$ | minimum contact energy between cells | 12 AUE/[lattice site length] |
| $J'_\alpha$ | contact energy between the cells and the medium | 24 AUE/[lattice site length] |
| $\nu$ | number of ligands and receptors per cell | 8 ligands–receptors |
| **Cell migration** | | |
| $\mu_{\text{pers}}$ | strength of persistent migration | 3 AUE |
| $\mu_{\text{chem}}$ | strength of chemotaxis | 1 AUE |
| $k_\chi$ | scaling factor chemoattractant gradient | 0.05 mol/[lattice site length] |
| **Cell division** | | |
| $\eta_{\text{init}}$ | delay before the start of division | 400 time steps |
| $\eta_{\text{grow}}$ | time for dividing cell to grow $2 \times A_{\text{t}}$ and divide | $2 \times 10^4$ time steps |
| **Food and metabolism** | | |
| $\tau_{\text{eat}}$ | refraction period of consumption of food | 2 time steps |
| $\tau_{\text{met}}$ | metabolic period | 100 time steps |
| $\Gamma$ | maximum number of food units supported on $\Lambda_{\text{food}}$ | 524288 |
| **Evolution** | | |
| $g$ | period between GRN updates | 20 time steps |
| $m$ | mutation probability | 0.05 |
| $\sigma_{\text{mut}}$ | standard deviation of mutation size | 0.05 |
| $\zeta$ | death probability | $5 \times 10^{-6}$ |

Table 2: Model parameters.

#### Supplementary videos

##### Video S1 Strictly unicellular life cycle.

##### Video S2 Multicellular + unicellular propagules life cycle.

##### Video S3 Multicellular propagules life cycle.

##### Video S4 High-adhesion multicellular life cycle.

**Video S5 Strictly unicellular lineage in EC 6. Cells can be seen dying while trying to reach food due to the large distances between food patches in this environment.**

**Video S6 High-adhesion multicellular lineage in EC 1. All cells aggregate in a single multicellular cluster that struggles to divide and take advantage of the available food.**
