## Supplementary figures and images for "Reproduction emerges from ecological interactions at the onset of multicellularity"

### Video s1

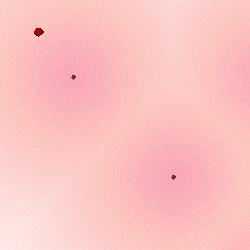

### Video S2

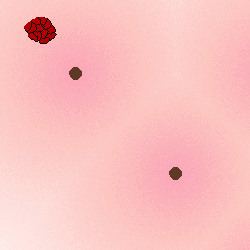

### Video S3

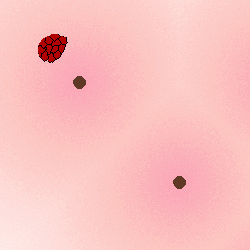

### Video S4

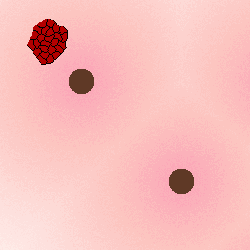

### Video S5

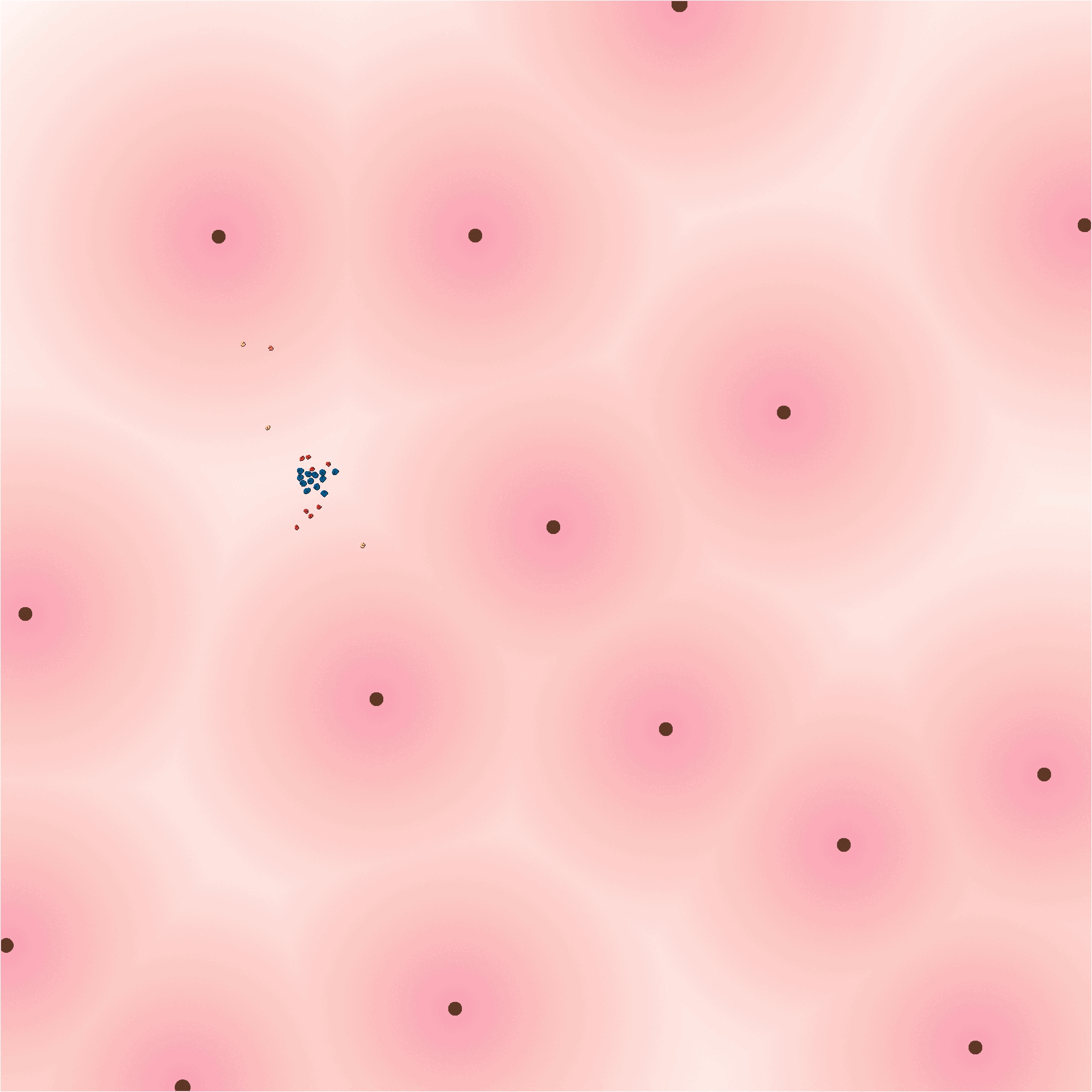

### Video S6

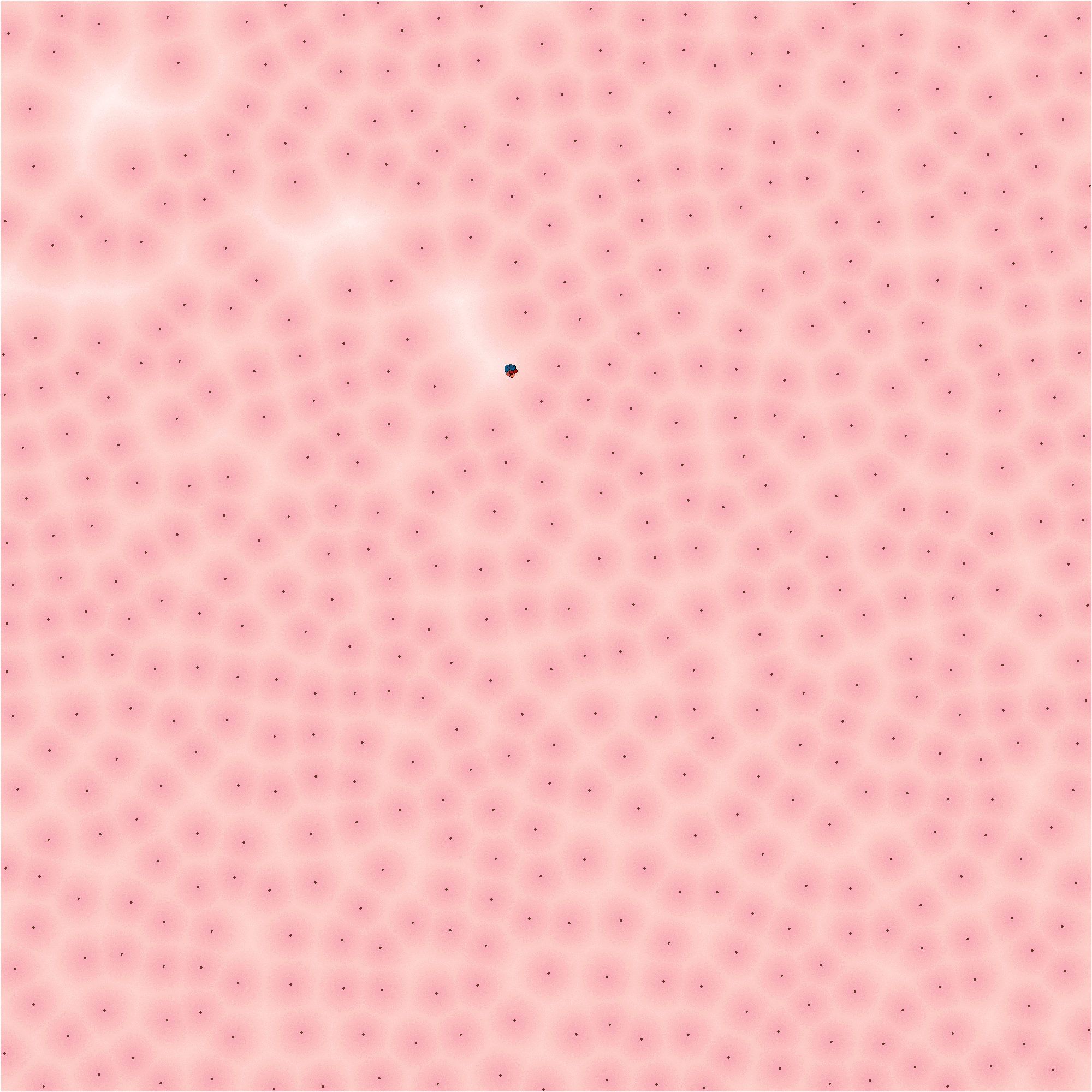
